# Heterotypic Condensate Formation by Prion Proteins and Pro-Inflammatory S100A9

**DOI:** 10.64898/2026.10.02.756240

**Authors:** Dominykas Veiveris, Viktorija Karalkevičiūtė, Kamilė Mikalauskaitė, Darius Šulskis, Mantas Žiaunys

## Abstract

Liquid-liquid phase separation (LLPS) of biomacromolecules has been shown to influence a large variety of biological processes via the formation of membraneless organelles. Aberrant LLPS, however, has been implicated in a number of oncological and neurodegenerative disorders. Proteins related to Alzheimer’s, Parkinson’s diseases and other major amyloidoses have all been shown to form these condensates. Interestingly, some amyloidogenic proteins were demonstrated to form heterotypic droplets, where both molecules coexisted within the same condensate and affected each other’s aggregation. While such cross-interactions have been reported numerous times under non-LLPS conditions, the heterotypic droplet assemblies were only shown for a very limited number of amyloidogenic proteins. In this work, we examine heterotypic LLPS of prion proteins and the pro-inflammatory S100A9, whose cross-interaction under non-LLPS conditions we confirmed in a previous work. The results show that this interaction under condensation-promoting conditions enhances their aggregation, leading to the formation of a higher concentration of amyloid structures. These findings suggest that the pro-inflammatory S100A9 may have a critical role in promoting prion protein aggregation via the assembly of heterotypic condensates under high molecular crowding conditions.

## Introduction

Protein aggregation in the form of amyloid fibrils is linked to a wide variety of amyloidoses, including the neurodegenerative Alzheimer’s and Parkinson’s diseases (1–3). Several decades of intense research into the topic of protein fibril formation have led to a general understanding of the mechanisms governing this structural transition (4–6). This new insight into amyloidoses has also yielded the discovery of several disease-altering modalities (7), such as Tafamidis for transthyretin amyloidosis (8) or Aducanumab for Alzheimer’s disease (9). However, the complex process of protein aggregation and gaps in knowledge still limit our ability to prevent or control the majority of amyloid-related disorders (7,10,11).

One factor that plays a major role in protein aggregation is the relatively recently discovered phenomenon of liquid-liquid phase separation (LLPS) (12,13). While the formation of membraneless organelles via LLPS is generally considered a natural process that is vital to many biological functions (12,14–16), uncontrolled aberrant LLPS has been implicated in various disorders (17,18), including amyloid-related neurodegenerative diseases (19,20). Over the last few years, it has been shown that several amyloidogenic proteins can form such high concentration condensates, which then undergo a liquid-solid transition. Aggregate formation within droplets was demonstrated for amyloid-beta (19), Tau (21), prion proteins (22), alpha-synuclein (23) and many others (24–27), suggesting that LLPS may be a critical step in amyloidogenic protein aggregation.

It was also recently discovered that a few amyloid-related proteins can form heterotypic condensates, where both molecules are present at high concentrations. While cross-interactions between different amyloidogenic proteins and their effect on aggregation have been reported numerous times (28–32), this LLPS-induced, high localized concentration of both proteins massively increases the incidence of potential interactions. Heterotypic droplet formation has been shown for several such pairings, including alpha-synuclein with Tau (33,34), S100A9 (35), TDP-43 (36) and prion proteins (37), as well as Tau with prion proteins (38). However, both the possibility of heterotypic droplet formation, as well as the resulting effect is still not explored for the majority of amyloid disease related protein pairings.

In our previous work, we discovered that prion protein 89-230 fragment (PrP 89-230) and the proinflammatory S100A9 can cross-interact *in vitro* and modulate the aggregation kinetics of PrP (39). We also observed that S100A9 can undergo both homotypic LLPS, which causes eventual S100A9 aggregation, as well as heterotypic LLPS with alpha-synuclein, which affects its droplet formation and aggregation kinetics (35). Additionally, according to the Protein Abundance Database (PAXdb) (40), S100A9 and prion proteins share a localization in multiple areas *in vivo*, including the cerebral cortex (41) and cerebrospinal fluid (42). Based on this information, there exists the possibility of heterotypic droplet formation by PrP and S100A9, where the high localized concentration would modulate the aggregation propensity of PrP.

Because the N-terminal part of prion proteins is known to modulate LLPS (43), we selected both full-length (23–230) and truncated (89–230) PrP variants for this study. To test the hypothesis of heterotypic condensate formation, we first examined PrP phase separation under a number of different environmental conditions, in an attempt to find ideal solution parameters for droplet formation *in vitro*. We then subjected S100A9 and PrP to these selected conditions and monitored their LLPS via brightfield and fluorescence microscopies, tracked real-time kinetics of liquid-liquid and liquid-solid transitions and analyzed the formed assembly secondary structures and morphologies. We observed that under LLPS-inducing conditions, the condensation and aggregation of this protein mixture is most prevalent under physiological pH conditions and an absence of denaturing agents, which goes against the usual highly denaturing conditions used for PrP aggregation. These findings suggest that heterotypic LLPS of PrP and the pro-inflammatory S100A9 may be a critical step in the onset and progression of prionopathies *in vivo*.

## Materials and Methods

### Protein purification

His-tagged mouse prion protein (89-230 and 23-230 variant) coding plasmids (pRESETB vector, Amp resistant) were transformed into E. coli BL21 (DE3) Star cells. The cells were grown in ZYM-5052 autoinduction media for 16 hours at 37°C and 220 RPM agitation. The purification of the prion proteins was conducted similar to a previously described protocol (44). In brief, the collected biomass was pelleted, resuspended in 6 M guanidinium hydrochloride (GuHCl), 100 mM potassium phosphate, 10 mM Tris-HCl and 10 mM glutathione buffer solution (pH 8.0) and sonicated. The cell lysate was centrifuged at 15 000 x g, after which the supernatant was filtered through a 0.22 µm pore-size filter and combined with 30 mL of IMAC Sepharose^TM^ 6 Fast Flow resin. After 30 minutes of incubation at 4°C with gentle stirring, the resin was loaded onto a column and connected to an Akta Start purification system. Protein renaturation was carried out on the column with a 200-minute gradient to 0 M GuHCl using the same buffer solution without the denaturant. Weakly bound proteins were washed out with a 50 mM imidazole, 100 mM potassium chloride, 10 mM Tris-HCl buffer solution (pH 8.0). The target protein was eluted using a 500 mM imidazole, 100 mM potassium phosphate and 10 mM Tris-HCl buffer solution (pH 5.8 for PrP 23-230, pH 6.4 for PrP 89-230). The collected protein was dialyzed in 4L of 10 mM sodium acetate (pH 4.0) buffer solution for 16 hours, concentrated to 200 µM (ε_280_ = 63 495 M^-1^cm^-1^ for PrP 23-230 and ε_280_ = 27 515 M^-1^cm^-1^ for PrP 89-230) and stored at -20°C.

S100A9 was purified as described previously (39). In brief, the plasmid containing 6xHis-SUMO-S100A9 was chemically transformed into One Shot^TM^ E. coli BL21 Star^TM^ (DE3) cells, that afterwards were grown at 37°C in LB medium containing kanamycin (50 mg/ml). Protein production was induced at OD_600_=0.7 with 0.4 mM isopropyl-β-d-thiogalactopyranoside and the cells were grown overnight at 25°C. Cells were pelleted, resuspended in 50 ml of lysis buffer (25 mM Hepes/NaOH, 0.5 M NaCl, and 10 mM imidazole (pH 7.5)) and sonicated. Cell debris was pelleted and the supernatant was applied to a Ni^2+^ Sepharose 6 Fast Flow (Cytiva) loaded gravity column, followed by stepwise elution with 20 ml of lysis buffer supplemented with 75 and 300 mM imidazole, respectively. 6xHis-SUMO-S100A9 protein fractions were dialyzed against 10 mM Tris buffer (pH 7.4) and the 6xHis-SUMO tag was cleaved with human Sentrin-specific protease 1 (SENP1) catalytic domain (derived from pET28a-HsSENP1, that was a gift from Jorge Eduardo Azevedo (Addgene plasmid #71465)) at 4°C overnight (45,46). The cleaved proteins were applied again to a Ni^2+^ column, and the flow-through was collected. The protein was further purified by size exclusion chromatography (HiLoad® 26/600 Superdex® 75 pg, Cytiva) in PBS, concentrated to 200 µM and stored at -20°C.

mCherry-S100A9 coding plasmid was prepared and the protein was purified as described previously (35), with immobilized metal affinity replacing anion exchange chromatography (47).

### Stock solution preparation

Polyethylene glycol (PEG, 20 kDa) stock solutions were prepared by dissolving 20 g of PEG in MilliQ H_2_O with the selected pH buffer components to yield final 40% PEG (w/v), 50 mL solutions with either 50 mM sodium acetate (pH 4.0 and pH 5.0) or 50 mM sodium phosphate (pH 6.0 and pH 7.0). To account for ionic strength differences of the solutions, each PEG buffer solution was supplemented with NaCl based on the Professor Robert J Beynon on-line buffer calculator [http://phbuffers.org/BuffferCalc/Buffer.html (accessed 2025-09-30)] to a final ionic strength of 200 mM.

The guanidinium hydrochloride (GuHCl) stock solutions were prepared by dissolving GuHCl to a final concentration of 7 M with either 50 mM sodium acetate (pH 4.0 and pH 5.0) or 50 mM sodium phosphate (pH 6.0 and pH 7.0). To account for the buffer component ionic strength differences (excluding GuHCl), the solutions were supplemented with NaCl as described previously.

Different pH buffer solutions without PEG or GuHCl were prepared with either 50 mM sodium acetate (for pH 4.0 and 5.0) or 50 mM sodium phosphate (for pH 6.0 and 7.0). The total ionic strength was adjusted to 200 mM as described previously.

The prepared stock solutions were stored at 4°C. Before use, they were thawed at 22°C for one hour with constant rotating mixing and filtered through a 0.22 µm pore size syringe filter.

### Brightfield and fluorescence microscopies

PrP stock solutions were combined with the PEG, GuHCl and other buffer solutions to yield a final protein concentration of 25 µM, 20% PEG and different pH (from 4 to 7), as well as GuHCl concentrations (0 M to 2 M). Samples containing S100A9 and/or PrP were prepared in 20% PEG, pH 7 solutions without GuHCl. The solutions were then incubated for 10 minutes at room temperature and 15 µL of each were placed on 1 mm-thick glass slides, covered with 0.18 mm coverslips. Brightfield images of the samples were acquired using an Olympus IX83 inverted microscope with a 40x objective (Olympus LUCPLANFL N 40x Long Working Distance Objective) as described previously (48). The image acquisition procedure was performed over a 10-minute time period after sample preparation. The resulting images were analyzed using Fiji ImageJ software (49). Only image brightness and contrast changes were applied during image analysis. Particle size and quantity was determined by fitting circular regions to the boundary of each visible protein assembly, calculating the radius and determining the theoretical volume of each particle (1 µm^3^ cut-off to exclude background anomalies).

Fluorescence and combined fluorescence-brightfield microscopy imaging was performed using 1 µM mCherry-S100A9 with or without 25 µM PrP (20% PEG, pH 7). Images were acquired using an Olympus IX85 inverted microscope with a 40x objective (Olympus LUCPLFLN40X2). Fluorescence and brightfield images were superimposed using Fiji ImageJ software (49). No brightness or contrast changes were applied during the analysis procedure.

### Optical density and fluorescence intensity

PrP solutions were prepared as described in the previous section and additionally supplemented with 100 µM thioflavin-T (from a 10 mM thioflavin-T stock solution). The samples were divided into 96-well Greiner F-bottom low protein binding plates, with 3 repeats for each condition and 200 µL volume in each well (the total preparation time was 30 minutes). The plates were placed in ClarioStar Plus plate readers with no cover at a set 22°C temperature. The optical density of each sample was scanned at 600 nm. The fluorescence intensity of ThT was acquired using 440 nm excitation and 480 nm emission wavelengths. After the data acquisition, the plate was covered with Nunc sealing tape and incubated at a constant 37°C temperature under quiescent conditions in the plate reader.

### Transmission electron microscopy (TEM)

After 3 day incubation at 37°C (20% PEG, different GuHCl concentrations and pH), the PrP samples were scanned using TEM as described previously (50). In brief, before sample application, 300 mesh formvar/carbon coated copper grids were glow discharged. 5 μL of PrP aggregate solutions were placed on the grids for 1 min, followed by drying with filter paper. Due to the presence of PEG, the grids were washed three times with 5 µL MilliQ H_2_O. The grids were negatively stained with 5 μL of 2% (w/v) uranyl acetate for 1 min, after which the excess solution was dried with filter paper. Finally, the grids were washed three times with 5 μL MilliQ H_2_O and dried with filter paper. All TEM images were recorded using Talos 120C (Thermo Fisher) transmission electron microscope, operating at 120V and equipped with 4k × 4k Ceta CMOS Camera. After acquisition, images were analyzed using Fiji ImageJ software (49).

### LLPS and aggregation kinetics

PrP and S100A9 stock solutions were combined with the PEG, GuHCl and other buffer stock solutions to yield reaction mixtures containing 25 µM PrP and 25 µM S100A9 (together and separately), 20% PEG, 0 or 0.5 M GuHCl and 100 µM ThT (final pH either 6 or 7). The solutions were then distributed to 96-well Greiner F-bottom low protein binding plates (3 repeats, 200 µL in each well) and the plates were covered with Nunc sealing tape. The plates were placed in ClarioStar plus plate readers and incubated at 37°C under quiescent conditions. ThT measurements were carried out every 10 minutes using 440 excitation and 480 nm emission wavelengths (0.1s resting time between each well scan). Kinetic data analysis was performed using Origin software.

### Fourier-transform infrared spectroscopy

After the LLPS and aggregation kinetic monitoring, each repeat of every sample was recovered and combined according to their reaction solution conditions. 500 µL of each reaction solution was centrifuged at 14 000 x g for 20 min, after which the supernatant was carefully removed. The aggregate pellets were then resuspended in 500 µL of D_2_O, supplemented with 400 mM NaCl, to improve sedimentation (51). Since the initial samples contained a high concentration of PEG, the centrifugation and resuspension procedure was repeated 4 times. After the final centrifugation, the supernatant was completely removed, and the aggregate pellets were resuspended in 50 µL D_2_O with 400 mM NaCl.

The FTIR spectra were acquired as described previously (51) using a Bruker Invenio S FTIR spectrometer. The acquired sample spectra were corrected by subtracting D_2_O and water vapour spectra, after which they were baseline-corrected and normalized in the 1595–1700 cm^-1^ range. Data analysis was performed using GRAMS software.

## Results

In order to test the hypothesis whether S100A9 and prion proteins (PrP) are capable of forming heterotypic droplets and how this interaction influences their subsequent liquid-solid phase transition, it was first necessary to determine optimal reaction conditions for both proteins. During our previous studies on S100A9 (35), we observed that it could form heterotypic condensates with alpha-synuclein under physiological pH conditions (PBS, pH 7.4) in the presence of a commonly used molecular crowding agent – polyethylene glycol (PEG). It could also form stable aggregates up to a ∼1.5 M concentration of a denaturant – guanidinium hydrochloride (GuHCl) (39). In the case of prion proteins, they have been shown to undergo both homotypic and heterotypic condensate formation under a number of different buffer pH, protein concentration, buffer component and additive conditions (37,38,52). Their aggregation into amyloid fibrils typically requires destabilizing additives, such as the aforementioned GuHCl (44).

Due to the lack of defined ideal conditions for PrP condensate formation, we first conducted brightfield microscopy imaging of truncated (89–230) and full length (23–230) mouse prion proteins under different pH and denaturant concentrations in the presence of 20% PEG (Supplementary Figures S1, S2). All of these factors – the N-terminal section of PrP (53), protein charge and folding (54)) – have been shown to modulate phase separation and/or amyloid aggregation.

In the case of the 23-230 PrP variant, we observed that large amounts of droplets formed under higher pH values and lower denaturant concentrations (Figure 1A, D). Droplet fluidity was confirmed by detecting their fusion events (Supplementary Figure S3). Smaller numbers of condensates were also visible at lower pH values and higher GuHCl concentrations (Figure 1A). Interestingly, when the solution was more acidic (pH 4.0 and 5.0), higher denaturant concentration conditions resulted in an almost immediate formation of PrP aggregates (Figure 1A, E). These results correlate with the sample optical density values (Figure 1B), where droplets and aggregates both caused notable increases in sample cloudiness, which could even be observed by simple visual inspection. The aggregates also possessed the ability to increase thioflavin-T (ThT, an amyloid-specific dye molecule) fluorescence intensity (Figure 1C), which can be caused by either the formation of amyloid fibrils or dye incorporation into rigid aggregate clusters. In the case of ThT fluorescence and the possible effect of pH on its signal intensity, the data was normalized by the intensity of ThT samples in the absence of PrP under their respective pH conditions.

**Figure 1.**
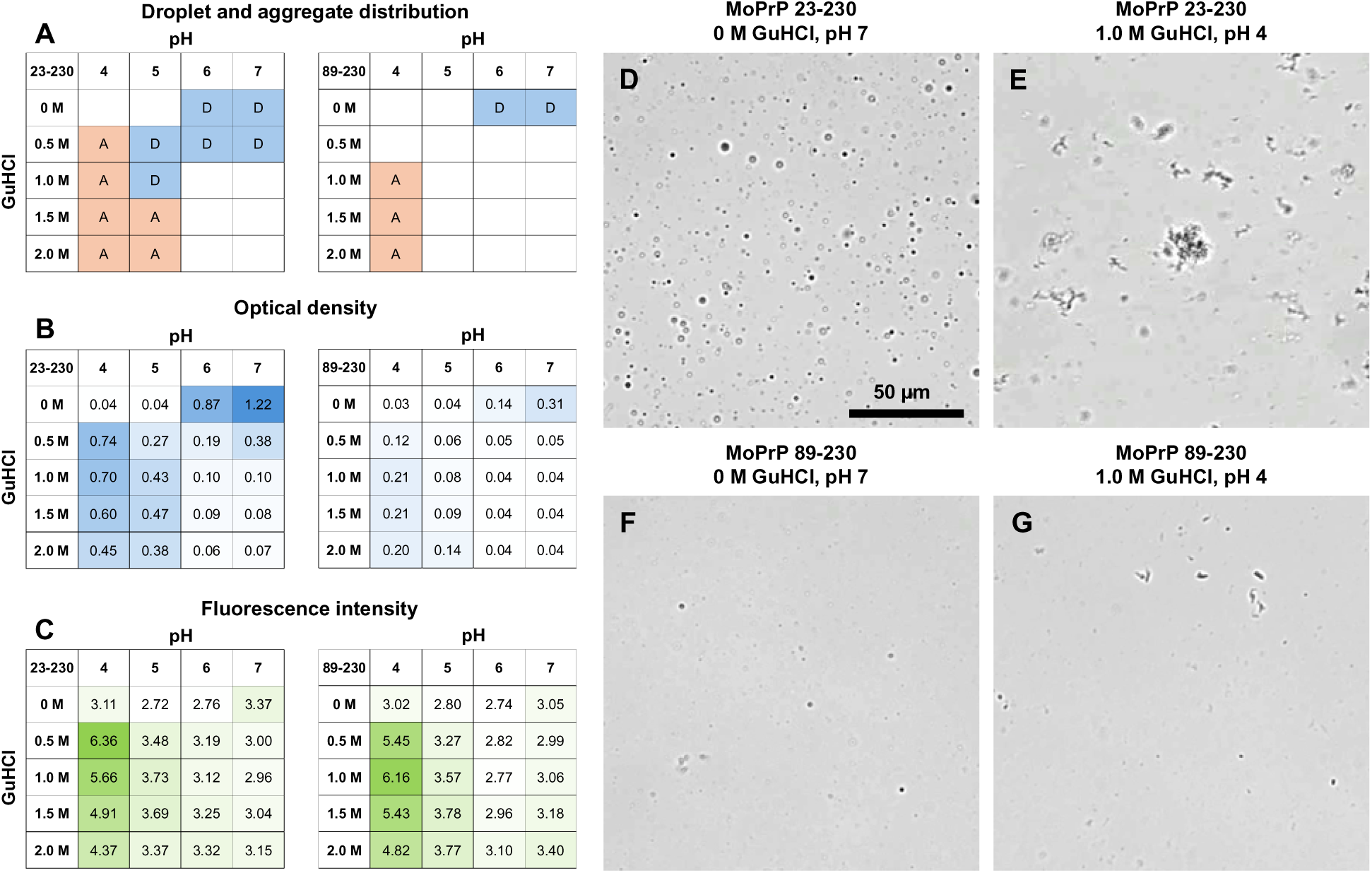
Liquid-liquid phase separation of full-length and truncated PrP under different environmental conditions. Samples that contain droplets or aggregates after 10 minutes of incubation under different pH and GuHCl concentrations (A, blue color – droplets, orange color - aggregates). The optical density (B) and ThT fluorescence intensity (C) of PrP samples under different pH and GuHCl concentrations (color gradient represents signal values from lowest to highest, each data point is the average of 8 technical repeats). ThT fluorescence was normalized by dividing the data by control ThT sample intensities (100 µM ThT in their respective pH buffer solutions without GuHCl).

When the truncated form of PrP (89–230) was subjected to the same range of conditions, there was a clear difference from the full-length variant. The number of visible droplets was massively reduced, and they appeared in a much more limited span of conditions (Figure 1A, F, G) and the same was true for the aggregates at lower pH values. This was reflected in the optical density data, where the values were smaller than in the case of the full-length PrP (Figure 1B). Interestingly, the fluorescence intensity of ThT within the samples that contained aggregates was similar for both protein variants (Figure 1C), despite clear differences in optical density (Figure 1B). This may indicate that only a part of the rapidly formed assemblies at low pH values contained ThT-positive structures.

During this assay, it was concluded that truncated PrP was considerably less suitable for LLPS monitoring due to its lower tendency for condensation under all of the examined solution variants. The full-length PrP, on the other hand, formed considerable numbers of condensates under several pH and GuHCl concentrations, making it possible to study its cross-interactions with S100A9 under multiple conditions (Figure 1A).

Brightfield images of 23-230 PrP under pH 7 and 0 M GuHCl (D), 23-230 PrP under pH 4 and 1.0 M GuHCl (E), 89-230 PrP under pH 7 and 0 M GuHCl (F), 89-230 PrP under pH 4 and 1.0 M GuHCl conditions (G, scale bar is 50 µm for all images).

To select the best conditions for the cross-interaction conditions, the full-length PrP samples that displayed droplet or aggregate formation were incubated for three days under quiescent conditions for 3 days at 37°C. The samples were then scanned using transmission electron microscopy (TEM) to determine which conditions resulted in the formation of fibril-like structures (Figure 2). Analysis by TEM revealed that all the samples contained fairly large structures, where aggregates were associated into large clusters. This is a common and expected occurrence with PrP aggregation, as the formed aggregate structures (both fibrillar and amorphous) have a high tendency to associate into sizeable clumps (55).

**Figure 2.**
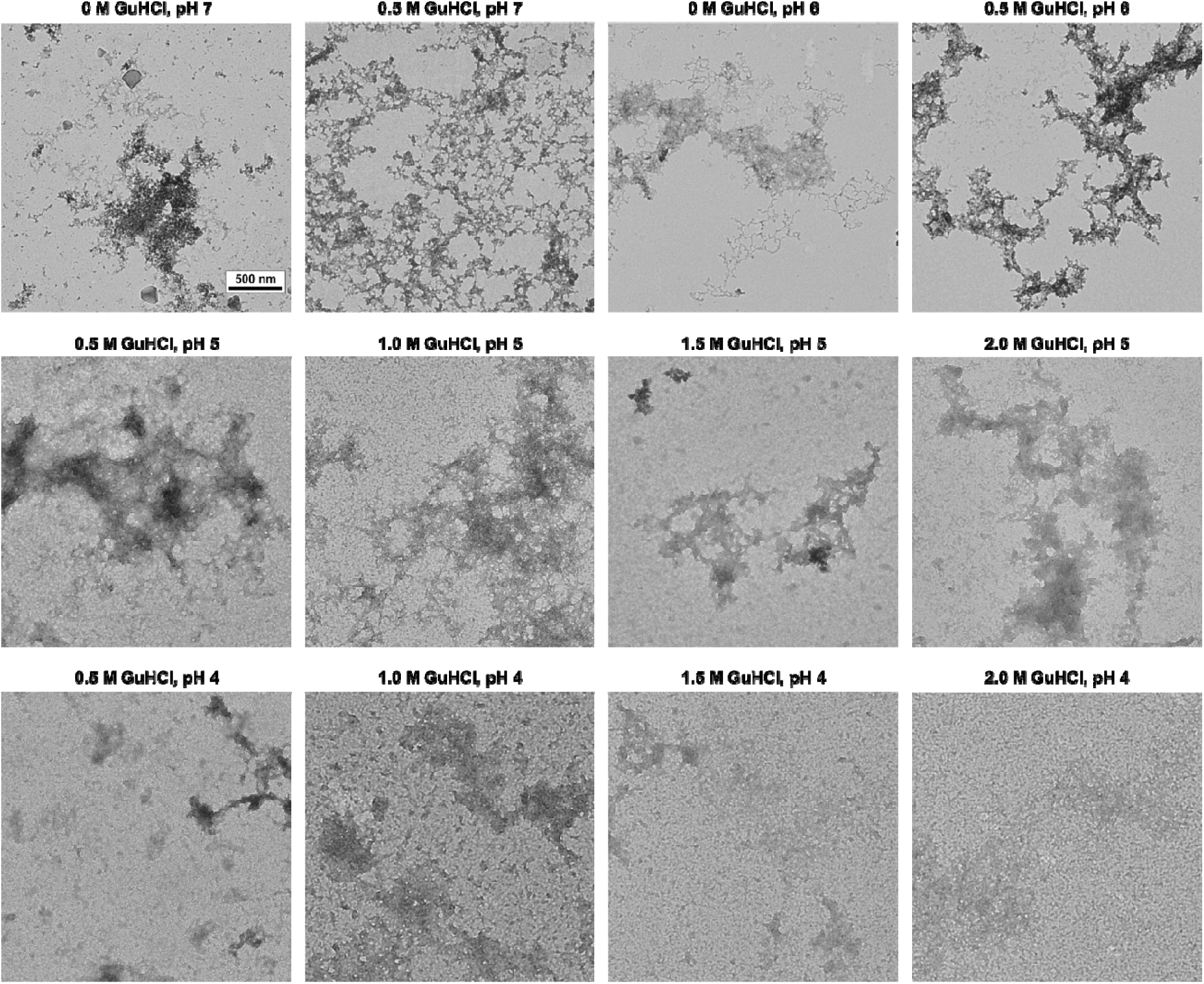
Transmission electron microscopy (TEM) images of full-length PrP aggregates under different environmental conditions. Images were acquired after each sample was incubated for 3 days under quiescent conditions at 37°C (scale bar is 500 nm for all images). Additional images of each sample are available as Supplementary Data.

Fibril-like structures were detected in pH 7 and pH 6 reaction solutions, which contained either 0 or 0.5 M GuHCl, with the most notable fibrillar assemblies seen in the absence of the denaturant. There also appeared to be a pH-dependent formation of aggregate clusters, where higher pH conditions contained mixtures of clumps and fibril-like species, while lower pH solutions contained mostly large clusters. Interestingly, LLPS-promoting conditions appeared to cause fibril-like structure formation at low denaturant concentrations, while non-LLPS conditions usually require strongly denaturing conditions. In addition to the protein destabilizing effects, higher GuHCl concentrations may influence droplet and aggregate formation due to its contribution to the solution’s total ionic strength.

Taking into consideration the results of brightfield microscopy and TEM imaging, four conditions were selected for PrP and S100A9 cross-interaction assays: pH 6 and pH 7 with 0 M or 0.5 M GuHCl in the presence of 20% PEG. Equimolar concentrations of both proteins (25 µM each) were subjected to these selected conditions, and their phase transitions were monitored by scanning the fluorescence intensity of thioflavin-T (ThT). The control PrP samples displayed two types of kinetic curves (Figure 3A-D, black curve). In two cases they were double-sigmoidal (pH 6, 0.5 M GuHCl and pH 7, 0 M GuHCl), which indicated a two-step process, where ThT fluorescence intensity first increases due to droplet formation and the dye’s localization within this crowded protein environment (56), followed by the second increase due to aggregate formation. Under the other two conditions (pH 6, 0 M GuHCl and pH 7, 0.5 M GuHCl), the ThT intensity increased very gradually throughout the entire incubation period with no clearly distinctive separation between LLPS and aggregation. Similarly, the S100A9 control samples also all displayed a gradual ThT intensity increase throughout the entire period (Figure 3A-D, red curve).

**Figure 3.**
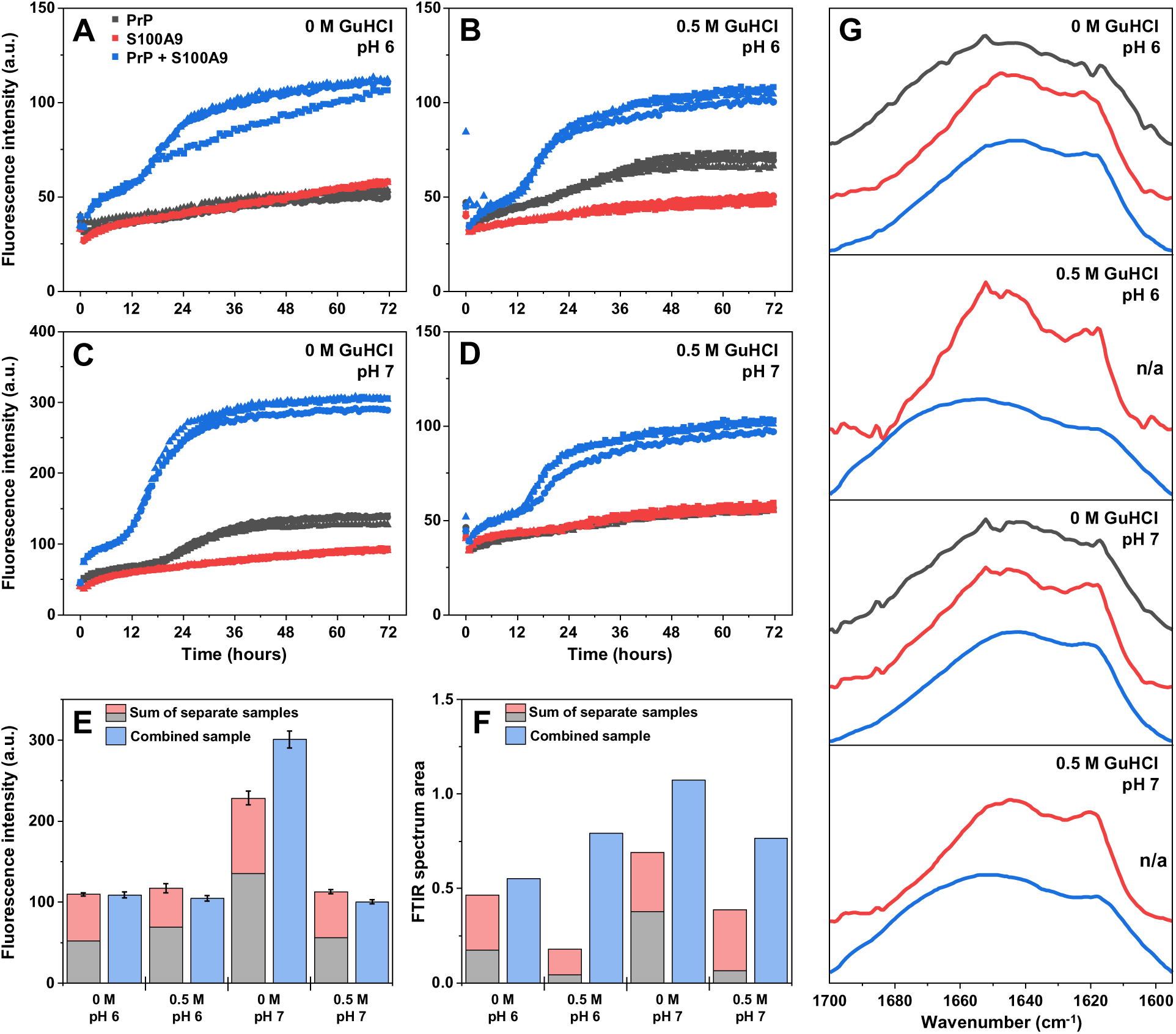
LLPS and aggregation of PrP and S100A9 under different pH and GuHCl concentration conditions. Aggregation kinetics of PrP and S100A9 monitored via ThT fluorescence intensity under pH 6 buffer solution with 0 M GuHCl (A), pH 6 with 0.5 M GuHCl (B), pH 7 with 0 M GuHCl (C), pH 7 with 0.5 M GuHCl (D) conditions. Sum of separate control sample and combined sample ThT fluorescence intensities (E, n=3, error bars are for one standard deviation) and FTIR spectra Amide I’ areas prior to normalization (F). Normalized FTIR spectra of PrP control (black curves), S100A9 control (red curves) and combined samples (G, blue curves). In cases where the sample did not contain aggregates after resuspension in D_2_O are marked as n/a.

When both proteins were present in the reaction solution, the kinetic curves all possessed a double-sigmoidal shape (Figure 3A-D, blue curve). The initial increase (droplet formation) occurred up to the 12-hour mark, which was followed by a much more pronounced shift in intensity, before reaching a plateau at the 24-hour mark. Comparing the end-point ThT fluorescence intensity revealed that in three of the four conditions, the sum of control sample intensities was similar to the mixture of both proteins (Figure 3E). Interestingly, the PrP and S100A9 mixture under pH 7 and 0 M GuHCl conditions displayed a higher ThT fluorescence intensity than the sum of both controls, indicating the formation of either additional ThT-positive or distinct structure aggregates (Figure 3E).

To compare the secondary structures of the formed aggregates, samples from each condition were pooled together and exchanged into D_2_O for FTIR analysis. During the centrifugation and resuspension steps, it was observed that the amount of aggregates formed in PrP control samples became increasingly lower with every resuspension step.

In the case of pH 6 buffer solution with 0.5 M GuHCl, as well as pH 7 with 0.5 M GuHCl, the PrP aggregates completely dissolved and obtaining their FTIR spectra became impossible (Figure 3G, black curves). In the other two conditions without GuHCl, the remaining precipitates displayed FTIR spectra with no clear peak related to beta-sheet content (∼1620 cm^-1^) (57). The integrated area of the Amide I’ region (1700 – 1595 cm^-1^) was also very low, which contributed to the high level of background noise in the spectra. These results suggested that the aggregates formed by PrP under LLPS conditions were either largely amorphous precipitates or their stability relied on the presence of 20% PEG.

In the case of S100A9 control samples, conversely to the PrP control, the aggregates remained stable even after multiple rounds of resuspension into D_2_O. The resulting spectra from pH 6 with 0 M GuHCl, as well as both pH 7 conditions were almost identical, with parallel beta-sheet-related peaks at 1620 cm^-1^, small anti-parallel peaks at 1695 cm^-1^ and broad peaks related to random-coil or alpha-helical regions at ∼1650 cm^-1^ (Figure 3G, red curves). They were also similar to S100A9 aggregates formed under non-LLPS conditions observed in previous studies (39). The only exception was the pH 6 with 0.5 M GuHCl condition S100A9 spectrum, which had a relatively larger peak at 1650 cm^-1^, suggesting the presence of more random-coil regions or amorphous aggregates.

When both proteins were in solution, the resulting assemblies were far more stable against dissolution during the resuspension procedure. This is most evident when comparing the sum of the control sample Amide I’ spectra area with the area of the combined sample (Figure 3F). Since the final aggregates were resuspended into an equal volume, this parameter is directly related to the number of protein amide bonds within each sample. In all four cases, the sum of control samples was lower than the area of the combined Amide I’ spectra. This difference was the highest in the pH 6 with 0.5 M GuHCl sample, where the control PrP became dissolved upon resuspension in D_2_O. The difference was also large in the case of both pH 7 conditions, where the pH 7 with 0 M GuHCl condition mixture had the largest Amide I’ area among all four samples.

The FTIR spectra of the combined PrP and S100A9 samples also had considerable differences in peak intensities (Figure 3G). Both pH conditions without GuHCl had a higher peak in the region associated with beta-sheet hydrogen bonding (∼1620 cm^-1^), while both conditions with 0.5 M GuHCl resulted in a reduced beta-sheet peak and larger random-coil and turn/loop regions (Supplementary Figure S4). This indicates that incubation of both proteins with the denaturant under LLPS-inducing conditions causes the formation of more amorphous structures that are stable to resuspension into D_2_O, or that the generated aggregates have larger random-coil, turn/loop regions.

Taking into consideration all the results, it appears that under high molecular crowding conditions, PrP and S100A9 undergo both LLPS and aggregation into beta-sheet rich structures under pH 7 in the absence of denaturants. These conditions result in the highest ThT fluorescence intensity changes, largest area FTIR Amide I’ spectrum and considerable beta-sheet content. While this is standard for the formation of S100A9 fibrils under non-LLPS conditions, it strongly contrasts with the high denaturation and agitation typically employed to force PrP aggregation. Based on all this data, pH 7 without GuHCl was chosen to monitor the possible heterotypic droplet formation by full-length PrP and S100A9.

When 25 µM of S100A9 was in the absence of full-length PrP under LLPS inducing conditions (20% PEG, 50 mM sodium phosphate, pH 7), brightfield microscopy images revealed that the protein formed a small number of tiny assemblies (Figure 4A, D-F, ∼500 detected particles from 3 images). Based on their distribution by volume and count, the dominant particles were between 1 and 2 µm^3^ in size. The 25 µM PrP sample displayed more than twice as many detectable particles under the same conditions (Figure 4B, D-F). They were also notably larger, with the dominant assemblies by volume being between 5 and 10 µm^3^. Unlike the control S100A9 sample, their distribution by count was fairly similar among the 1 – 2, 2 – 5 and 5 – 10 µm^3^ groups. When both proteins were present in solution, the volume and count distributions were similar to the prion protein control sample (Figure 4C, D-F), however, the number of detectable particles was much greater than the sum of both control samples (Figure 4F).

**Figure 4.**
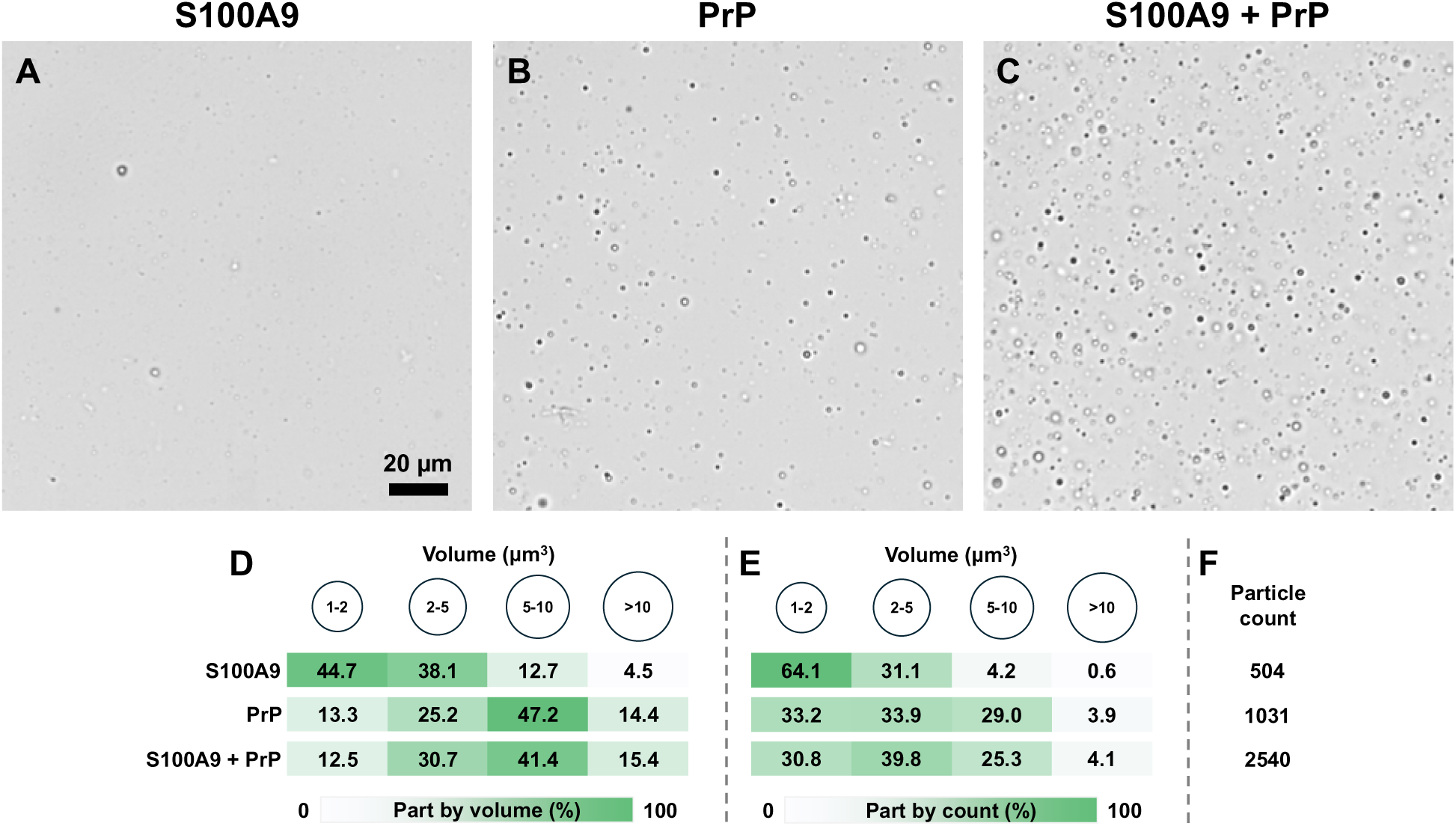
Condensation of S100A9 and full-length PrP under LLPS-inducing conditions. Brightfield images of control 25 µM S100A9 (A), 25 µM PrP (B) and S100A9 with PrP (C). Distribution of different particles by their total volume (D) and count (E). Number of detected particles from three separate images (F). Additional brightfield images are available as Supplementary Data (scale bar is 20 µm for all images).

If the combined sample displayed an intermediate particle size distribution between the control S100A9 and PrP samples, it would indicate that the proteins did not cross-interact and the rise in particle count was due to an increase in total protein content. However, the combined sample appeared to contain assemblies that were similar in size to the PrP control, but in a much greater quantity. This would suggest that the presence of both proteins promoted condensation and S100A9 may have been incorporated into the PrP droplets. In order to test this hypothesis, 1 µM of mCherry-S100A9 was combined with 25 µM PrP under the same LLPS-inducing conditions.

When mCherry-S100A9 was present in solution without PrP, it formed a small number of tiny visible droplets (Figure 5A). We have previously observed this effect, where the addition of a fluorescent tag increased the LLPS potential of S100A9 (35). When it was combined with 25 µM PrP, both the number and size of detectable assemblies increased, indicating that the labeled S100A9 became incorporated into condensates formed by PrP (Figure 5B). Analysis of the protein mixtures via combined brightfield and fluorescence microscopy revealed an interesting distribution of the labeled protein. It was observed that only part of the droplets contained mCherry-S100A9 and the relative amount present in each condensate was different based on their fluorescence intensity. These results suggest PrP can form both homotypic, as well as heterotypic droplets within the same protein mixture. A similar phenomenon was observed during our previous study with alpha-synuclein, where S100A9 was incorporated into only a part of the condensates (35).

**Figure 5.**
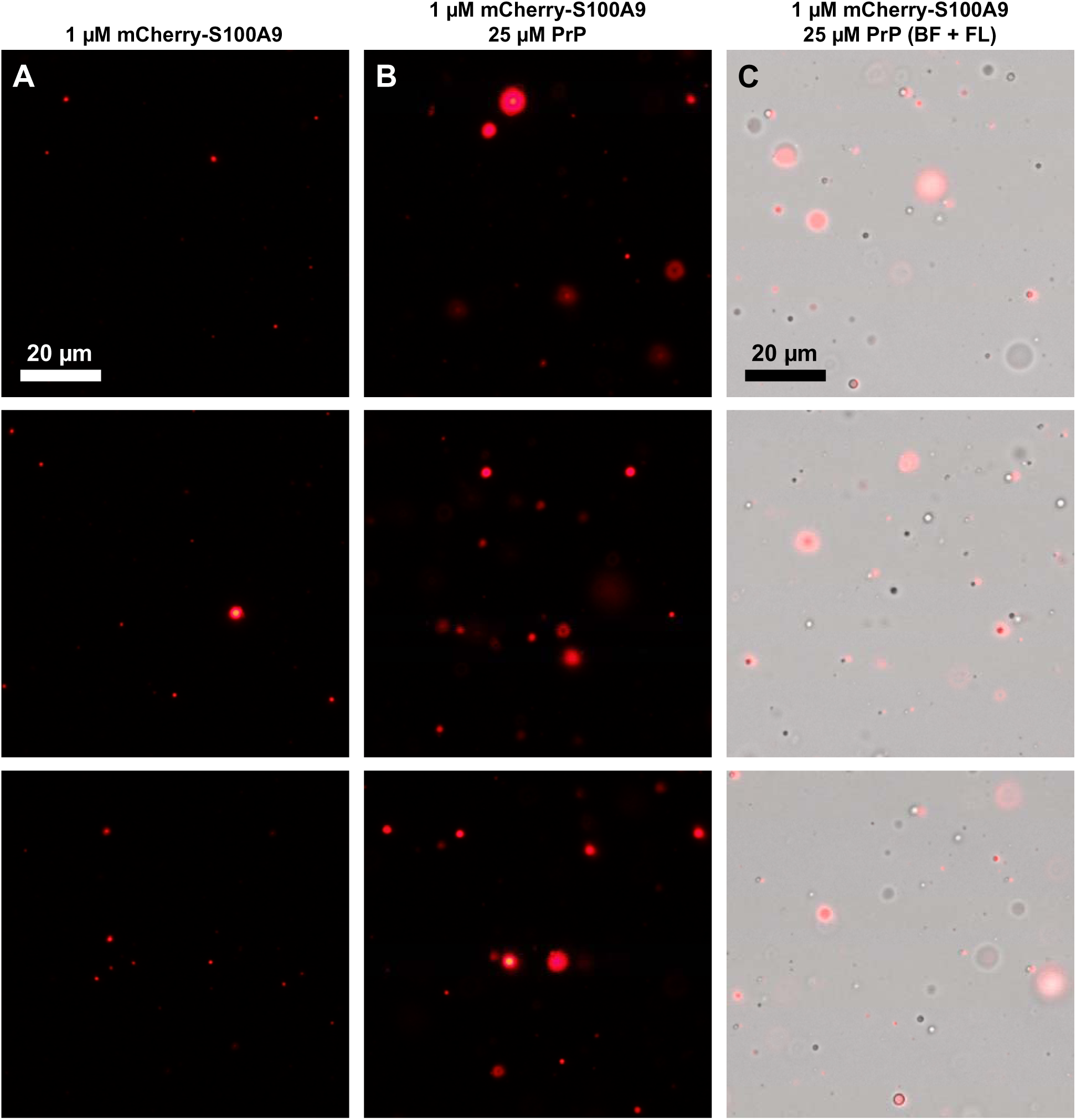
Heterotypic condensate formation by mCherry-S100A9 and PrP. Fluorescence microscopy images of 1 µM mCherry-S100A9 in the absence (A) and presence (B) of 25 µM PrP. Combined brightfield and fluorescence microscopy images of 1 µM mCherry-S100A9 in the presence of 25 µM PrP (C). Scale bar is 20 µm for all images.

## Discussion

In this study, we investigated the possibility of heterotypic LLPS between the pro-inflammatory S100A9 and prion proteins under different environmental conditions. Alongside the confirmation that these proteins can cross-interact during condensation, which affects their aggregation, we also observed a peculiar effect that LLPS-inducing conditions have on the demixing and aggregation of PrP.

During the initial condition optimization experiments, the LLPS-inducing environment significantly impacted the stability and behaviour of PrP. Low pH conditions, which are typically used for purified recombinant PrP storage (44), resulted in an almost immediate PrP transition to its aggregated state. A small portion of these formed structures even had the capacity to bind ThT, resulting in an amyloid-specific increase in fluorescence intensity. In the case of neutral pH, where aggregate formation is usually promoted by employing strong denaturants (44), the highest concentration of condensates and incubation-induced fibril-like structures assembled in the absence or low concentrations of GuHCl. This disparity between non-LLPS and condensation-promoting environment effect on PrP allowed its cross-interaction with S100A9 to be examined under more physiologically relevant conditions.

During sample analysis via combined brightfield and fluorescence microscopy, it was observed that labeled S100A9 became incorporated into PrP droplets. However, the protein mixture appeared to contain both homotypic (composed of just PrP), as well as heterotypic condensates, which had varying levels of mCherry-S100A9. Statistical analysis of the droplet size and quantity distribution revealed that S100A9 does not simply become included into PrP condensates (reduced part of small assemblies) but also promotes the overall demixing process (increased number of detectable particles). Previous studies with heterotypic droplet formation by PrP and other amyloidogenic proteins have also shown the influence of such complexes on condensate characteristics. Alpha-synuclein was demonstrated to accelerate amyloid formation when present with PrP under LLPS-inducing conditions (37). In the case of PrP and Tau cross-interactions (38), the droplets became smaller and had lower fluidity.

Based on the observations in this study and previously published work on PrP complex formation with other proteins under LLPS-inducing conditions (37,38), the most likely factor that leads to heterotypic droplet formation of S100A9 and PrP is electrostatic interactions. Under conditions where the droplet formation and cross-aggregation is most evident (pH 6 – 7), PrP 23-230 has a positive charge (pI = 9.64), while S100A9 is charged negatively (pI = 5.72). This difference could facilitate both protein interactions not only under non-LLPS conditions, as observed previously (39), but also in a LLPS-inducing environment. The electrostatic interactions would lower the energy barrier for both protein demixing, as well as promote heterotypic condensate formation. The negative effect of an increased solution ionic strength due to the addition of GuHCl, which is best observed in the ThT intensity change during incubation and end-point FTIR spectra area (Figure 3), further supports this idea.

In the case of the onset of prionopathies, this cross-interaction and heterotypic condensate formation may be one of the underlying factors for the onset of PrP-related disorders. During this study, it was observed that the PrP aggregates, which form during homotypic LLPS, are meta-stable and dissolve upon multiple rounds of resuspension in a solution without PEG. However, the addition of S100A9 causes the formed aggregates to retain their stability even under non-LLPS conditions. Furthermore, these assemblies have a larger ThT binding capacity and result in a higher amyloid-specific fluorescence signal.

Taken together, these results demonstrate the potential influence of S100A9 on the condensation and aggregation process of PrP. This heterotypic condensation-driven formation of higher stability amyloid assemblies may be a potential pathway for the onset of prion-related disorders.

## Data availability

The raw data used for the preparation of this manuscript and additional transmission electron, bright field and fluorescence microscopy images are available at Mendeley Data: https://data.mendeley.com/datasets/wjmdsbxfwf/1

