## Supplementary information for "Heterotypic Condensate Formation by Prion Proteins and Pro-Inflammatory S100A9"

**Supplementary Material**

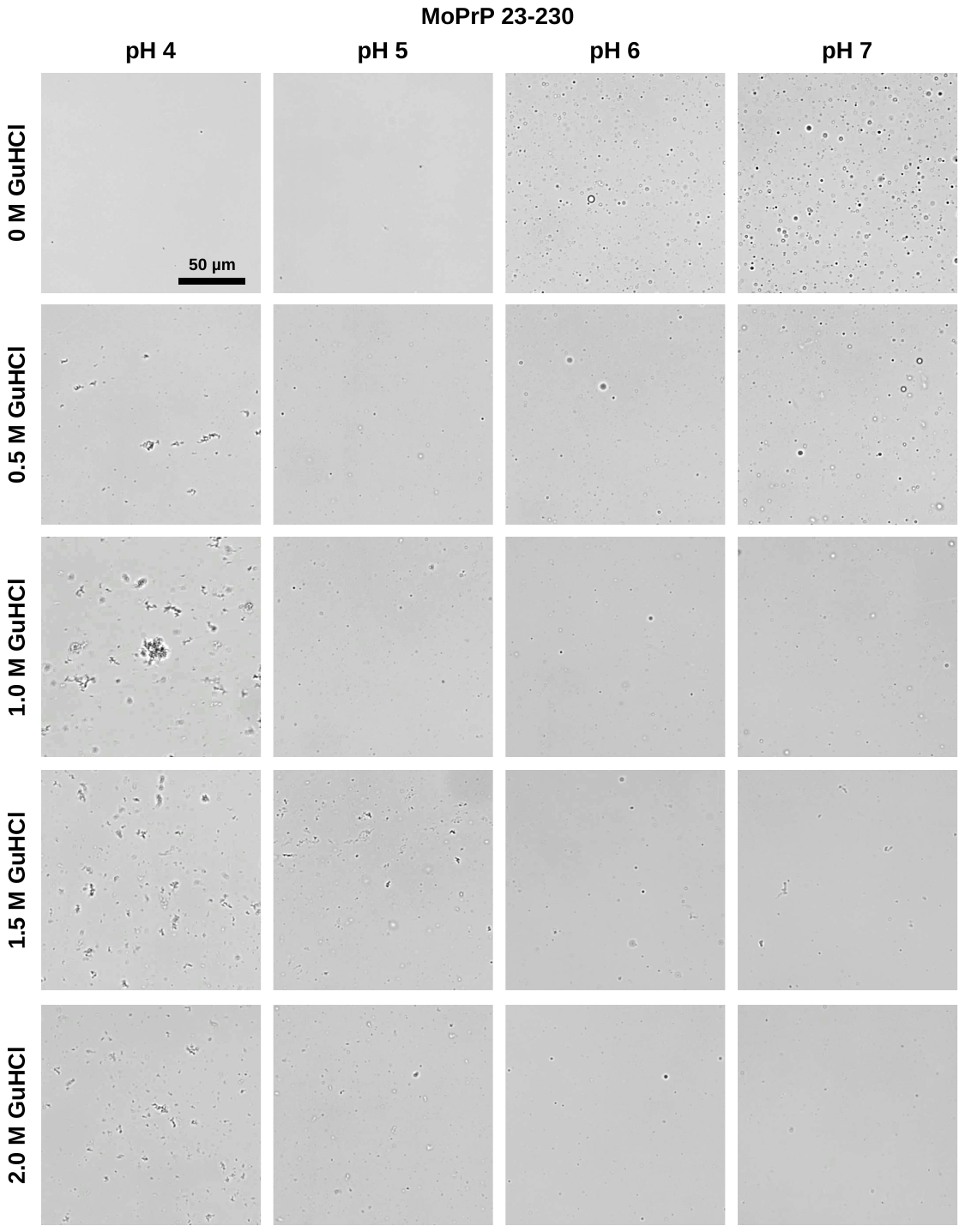

**Figure S1.** Brightfield microscopy images of 25 µM MoPrP 23-230 under different GuHCl concentrations and pH in the presence of 20% PEG.

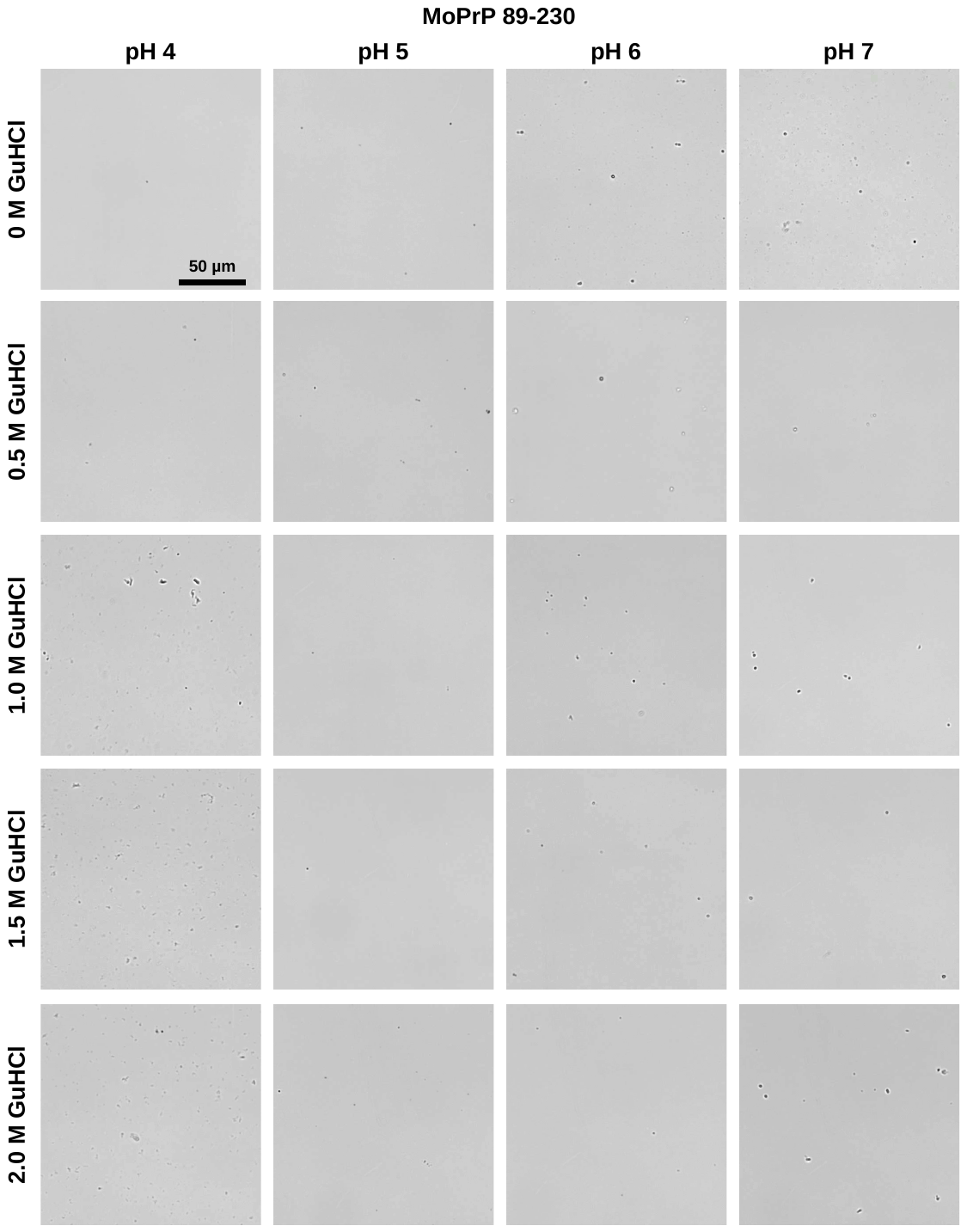

**Figure S2.** Brightfield microscopy images of 25 µM MoPrP 89-230 under different GuHCl concentrations and pH in the presence of 20% PEG.

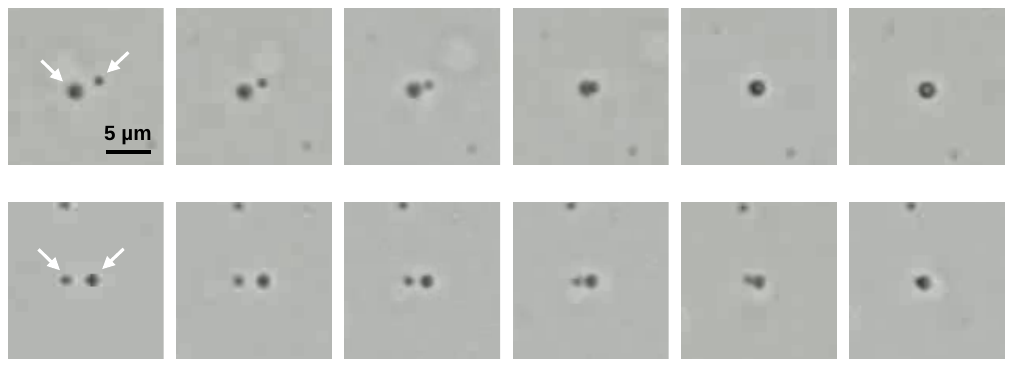

**Figure S3.** Fusion of PrP (23-230) droplets. 25 µM PrP, 20% PEG, 50 mM sodium phosphate (pH 7.0), 22°C. Scale bar is 5 µm.

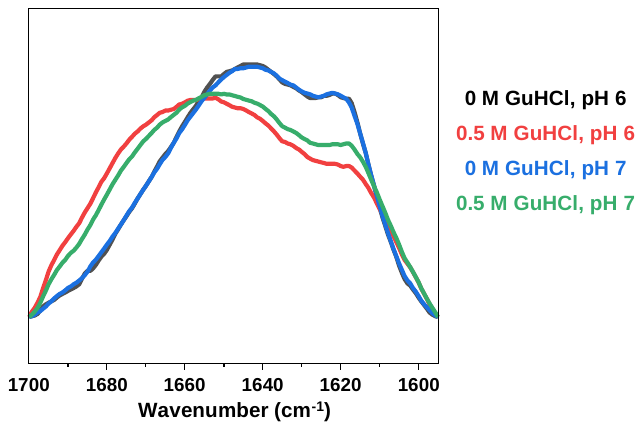

**Figure S4.** FTIR spectra comparison of combined 25 µM S100A9 and 25 µM PrP aggregates, formed after incubation under LLPS-inducing conditions.
